# NeuroTorch: A Python library for neuroscience-oriented machine learning

**DOI:** 10.1101/2024.12.29.630683

**Authors:** Jérémie Gince, Anthony Drouin, Patrick Desrosiers, Simon V. Hardy

## Abstract

Machine learning (ML) has become a powerful tool for data analysis, leading to significant advances in neuroscience research. While ML algorithms are proficient in general-purpose tasks, their highly technical nature often hinders their compatibility with the observed biological principles and constraints in the brain, thereby limiting their suitability for neuroscience applications. In this work, we introduce NeuroTorch, a comprehensive ML pipeline specifically designed to assist neuroscientists in leveraging ML techniques using biologically inspired neural network models. NeuroTorch enables the training of recurrent neural networks equipped with either spiking or firing-rate dynamics, incorporating additional biological constraints such as Dale’s law and synaptic excitatory-inhibitory balance. The pipeline offers various learning methods, including backpropagation through time and eligibility trace forward propagation, aiming to allow neuroscientists to effectively employ ML approaches. To evaluate the performance of NeuroTorch, we conducted experiments on well-established public datasets for classification tasks, namely MNIST, Fashion-MNIST, and Heidelberg. Notably, NeuroTorch achieved accuracies that replicated the results obtained using the Norse and SpyTorch packages. Additionally, we tested NeuroTorch on real neuronal activity data obtained through volumetric calcium imaging in larval zebrafish. On training sets representing 9.3 minutes of activity under darkflash stimuli from 522 neurons, the mean proportion of variance explained for the spiking and firing-rate neural network models, subject to Dale’s law, exceeded 0.97 and 0.96, respectively. Our analysis of networks trained on these datasets indicates that both Dale’s law and spiking dynamics have a beneficial impact on the resilience of network models when subjected to connection ablations. NeuroTorch provides an accessible and well-performing tool for neuroscientists, granting them access to state-of-the-art ML models used in the field without requiring in-depth expertise in computer science.

## 1 Introduction

The early stages of machine learning (ML) were deeply influenced by neuroscience.^1^ Key influences include the first abstract neuron models^2^, artificial neural networks for data classification^3^, recurrent neural networks with associative memory^4^, and reinforcement learning^5^. Today, ML is reciprocating this influence by driving significant progress in the field of neuroscience. It facilitates the analysis of vast amounts of data generated by brain imaging techniques such as functional magnetic resonance imaging (fMRI)^6^ and electroencephalography (EEG)^7^ as well as the segmentation of cells in microscopy data^8^. Machine learning also aids in the diagnosis of brain diseases, such as Alzheimer’s disease^9^, Parkinson’s disease^10^, and schizophrenia^11^, and contributes to the modeling of neural processes, including organizational principles in the sensory cortex^12^. Despite these advances, ML has yet to be broadly adopted across all neuroscience research. This may be attributed to the computational resources and advanced programming skills required for the implementation of some machine learning techniques, as well as a current lack of models that are either inspired by or constrained by biological plausibility, which would likely be more interpretable to neuroscientists.^1, 13, 14^

Among all Machine Learning (ML) techniques, Recurrent Neural Networks (RNNs) [Fig. 1.1 (a)] uniquely incorporate fundamental structural features inspired by biological neural networks^15, 16^, such as recurrent connections that mimic neural feedback loops and the capacity for both local and long-range connections. On the functional level, RNNs operate based on non-linear dynamics governed by threshold-like activation functions, echoing the dynamical behavior regulating the firing rates of interconnected biological neurons^17^. Furthermore, RNNs are recognized as universal approximators of dynamical systems^18, 19^, meaning that any time series can be modeled as the output of an adequately complex RNN. Consequently, RNNs exhibit a rich variety of dynamical behaviors - from attractor dynamics, through oscillatory, to even chaotic regimes. This diversity offers a versatile framework for understanding how computations can be performed by neural populations^20^.

**Figure 1.1.**
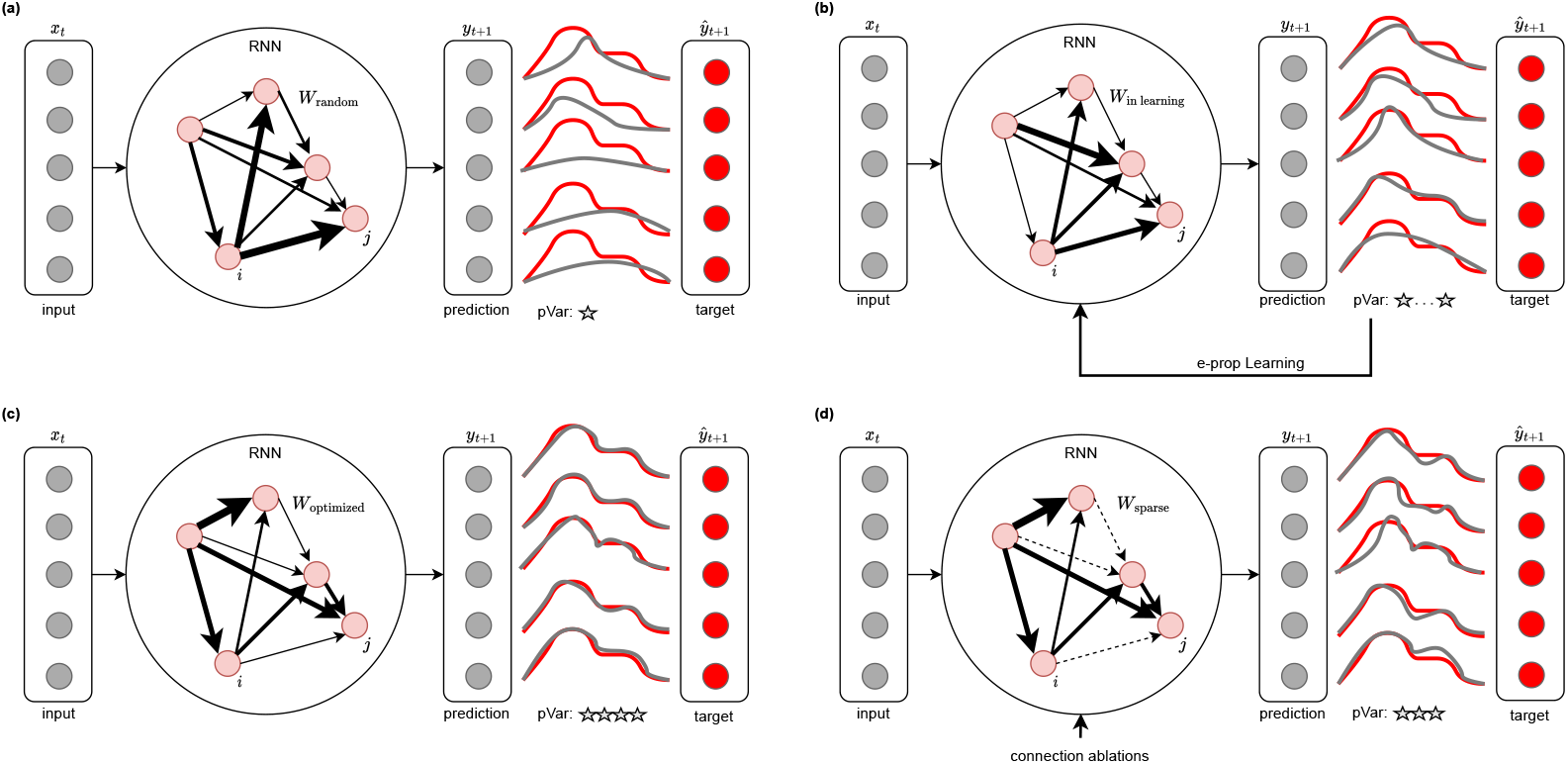
Numerical experiment during which a RNN is trained to fit time series data. (a) A recurrent model is created with random weights to make time series predictions. The gray traces represent the model’s predicted traces, while the red traces represent the targets. The pVar of these predictions is poor. (b) The model is then trained with the e-prop algorithm to optimize its weights and reproduce the target time series. The pVar gradually increases during training. (c) The now trained model is capable of reproducing the target time series with a high pVar. (d) We remove weights from the model to make it more sparse and observe the impact on the pVar.

Owing to a diverse range of training algorithms^21–24^, which especially permit the modification of connection weights to fit target data [Fig. 1.1 (b)–(c)], RNNs have garnered significant successes in neuroscience. They have been instrumental in unveiling altered intra-habenula interactions contributing to behavioral passivity in larval zebrafish^25^, and identifying single-cell targets for circuit control in epileptic animals^26^. Very recently, RNNs have demonstrated high precision in replicating neuronal recordings and forecasting the behavior of animals in decision-making or reward-learning tasks.^27–30^ The rapid progress in this field strongly suggests that single RNN models may soon possess the capability to simulate neural activity during complex behaviors and various perturbations within the relevant neural circuitry or behavioral contexts, making them a highly potent tool for systems neuroscience when utilized as surrogates for new experimental data.^31^

Another fundamental principle bridging ML and neurobiology comes from spiking neural networks (SNNs), which offer relatively realistic biological dynamics through their spike-based activity.^32, 33^ In addition, the training of those networks substitutes the traditional backpropagation training with biologically plausible processes such as Spike-Timing-Dependent Plasticity (STDP)^34^ and eligibility trace forward propagation (e-prop)^35^, using local information rather than global gradients^36, 37^. By adopting these biologically inspired dynamics and learning mechanisms, deep learning models can better align with real neural processes, enabling the study of biological phenomena through numerical simulations. The emergence of neuromorphic processors, such as Intel’s Loihi^38^ and SpiNNaker^39^, has played a crucial role in the growing popularity of spiking neural networks^40^. These processors leverage the specific characteristics of SNNs, including their sparsity and asynchronous nature, allowing for efficient time and energy utilization.^41–43^

All these advances plead in favour of systematically incorporating SNNs in any ML framework intended to assist neuroscientists in data analysis and neuronal mechanism modeling. However, the mainstream ML libraries like TensorFlow^44^ and PyTorch^45^, both written in Python^46^, do not natively support SNNs. These libraries are primarily focused on the traditional artificial neural networks, and they do not include out-of-the-box support for the special neuron models and learning rules that are characteristic of SNNs. Even high-level librairies built on top of PyTorch, such as PyTorch Lightning^47^ and Poutyne^48^, while providing invaluable tools to expedite and simplify PyTorch development processes without compromising flexibility and control, do not extend their offerings to include models or training methods drawn from neuroscience.

To address this gap, more specialized libraries like Norse^49^, SpyTorch^50^, and snnTorch^51^ have been developed to implement spiking neural network architectures, facilitating research in neuromorphic computing. These Python-based libraries provide a set of spiking learning layers that can be used with PyTorch, demonstrating their functionality in tasks such as the classification of popular datasets like MNIST^52, 53^ and Heidelberg^54^. A more extensive catalog of SNN-centric libraries, each offering a diverse assortment of learning rules, is presented in the paper by Manna et al.^55^. Despite the breadth of available options, none of these libraries currently incorporate the e-prop learning rule —an algorithm of significant relevance given its biological plausibility and superior performance as demonstrated by Bellec et al.^35^. In addition, other Python packages have successfully incorporated a certain number of neurobiological constraints and techniques. The package proposed by Suarez et al.^56^ constructs recurrent neural networks (RNNs) with firing-rate dynamics and network architecture based on real connectomes. These RNNs can be partially trained using reservoir computing, where only the weights of the readouts are trained, to evaluate their ability to store information. Although the learning rule is not biologically realistic and is limited to a portion of the network, this approach is relevant as it incorporates biological constraints through the network structure. Concurrently, yet another package has been developed by D’Amicelli et al.^57^, which also employs reservoir computing with a connectome-constrained structure.

Given the pressing need for more realistic neural network models, including biologically plausible learning algorithms, coupled with the current lack of comprehensive ML libraries offering learning pipelines specifically tailored to neuroscience research, we have developed NeuroTorch^58^. This library, built upon PyTorch, leverages the latest advances in RNN and SNN libraries. It provides features for neuronal time-series analysis, regularization based on connectome metrics, and optimization methods such as backpropagation-through-time (BPTT)^36, 59^, eligibility trace forward propagation (e-prop)^35^, recursive-least-square (RLS)^60–63^, and proximal policy optimization (PPO)^64^. The library also incorporates popular dynamical models for neuronal activity, such as the firing-rate dynamics of Wilson-Cowan (WC)^17, 65–68^ and the spiking dynamics leaky-integrate-and-fire with explicit synaptic current (SpyLIF)^50^, along with other tools for neuromorphic computing research.

The paper presents an in-depth exploration of NeuroTorch. We start in Section 2 by highlighting the results obtained from various numerical experiments, validating the performance of NeuroTorch. Specifically, we demonstrate that NeuroTorch achieves comparable results to other existing packages when tested on benchmark datasets such as MNIST, Fashion-MNIST, and Heidelberg. Furthermore, we delve into the reproduction of neuronal activity patterns in the ventral habenula of a larval zebrafish using NeuroTorch’s e-prop implementation. This showcases the capability of NeuroTorch to handle time series data from experimental measurements. Finally, we explore the concept of resilience in relation to sparsity and hierarchical ablation, shedding light on the robustness of NeuroTorch’s trained models [Fig. 1.1 (d)]. Moreover, we investigate the impact of Dale’s principle (a.k.a Dale’s law)^69, 70^ on the trained models, providing insights into the significance of incorporating realistic biological rules in enhancing model performance and resilience against connection ablations. Figure 1.1 gives a conceptual overview of our computationnal experiments. In Section 3, we dissect these results further and identify certain limitations in our work while also proposing future avenues for exploration. Finally, Section 4 offers a detailed description of NeuroTorch, outlining its design, implementation, and key features, to provide readers with essential technical information about the framework.

## 2 Results

This section presents the results of our computational experiments conducted with NeuroTorch^58^, the library developed and introduced in this paper. The experiments include testing the NeuroTorch pipeline with BPTT learning algorithm on benchmark datasets, demonstrating its effectiveness in classification models based on SNNs. Furthermore, we explore the application of NeuroTorch in the analysis of real time series data, specifically focusing on neuronal measurements of the ventral habenula in a larval zebrafish. By optimizing firing-rate and spiking dynamics using the e-prop learning algorithm, we assess the effectiveness of NeuroTorch in handling time series data and capturing the dynamics of neuronal activity. Additionally, we investigate the resilience of the optimized networks against connection loss and examine its consequences on the properties of the connectome produced by the type of neuronal dynamics employed, as well as the imposition of yet another biologically realistic rule, namely Dale’s Law. This analysis allows us to gain insights into the models’ ability, when submitted to biologically plausible constraints, to maintain performance and adaptability in the face of parameter perturbations or changes in their connectivity.

### 2.1 Data classification benchmark

Before dealing with typical neuroscience data, we evaluated the performance of the NeuroTorch package and its BPTT algorithm by conducting experiments on three commonly used benchmark datasets for classification, namely MNIST^52, 53^, Fashion-MNIST^71^, and Heidelberg^54^. The two former datasets contain grayscale images of handwritten digits and fashion products, respectively, while the latter dataset contains spike trains representing audio recordings of spoken digits in both German and English. Our objective was to assess the functionality of NeuroTorch and determine how it compares to other established packages (SpyTorch^50^ and Norse^49^) in terms of prediction accuracy when performing classification tasks.

The SNN models implemented using NeuroTorch, along with other selected packages, were tested to ensure fair and comparable evaluations. It is important to note that there are some differences in the training and implementation methods, which justify the variations in the results [see the method for more details on the implementation]. The models used in these experiments were not necessarily optimized specifically for the task at hand, but rather designed to provide a baseline assessment of the packages’ performance. The results presented in Table 2.1 demonstrate that NeuroTorch performs comparably to the other packages. The models implemented using NeuroTorch achieve prediction accuracies that are on par with the established packages, confirming that NeuroTorch functions as intended.

**Table 2.1.**
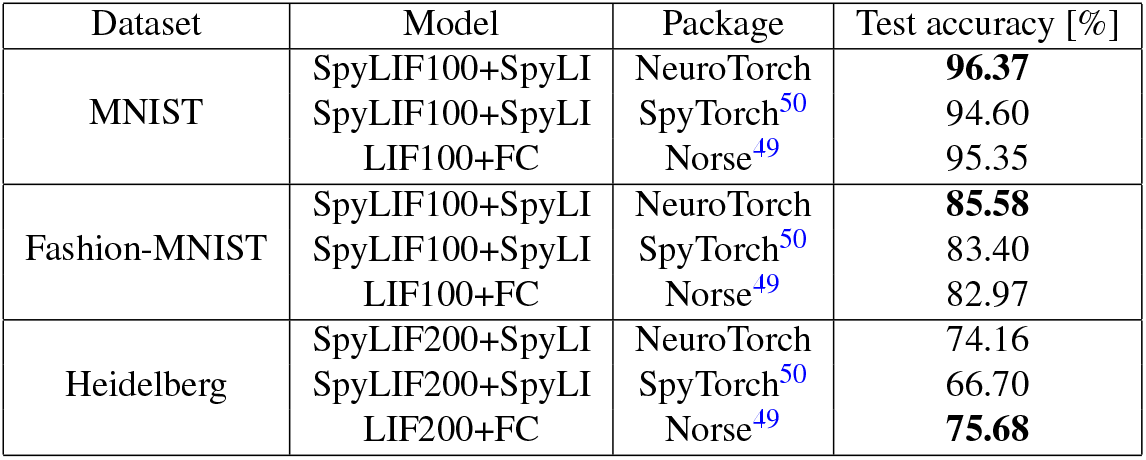
Prediction accuracy of different SNN classification models computed with the test-subset of the benchmark datasets MNIST, Fashion-MNIST and Heidelberg. The models are named as “<Input Dynamics><number of units>+<Output Dynamics>”. The spiking dynamics are defined in Section 4.1. The number highlighted in bold represents the highest achieved performance for each dataset, indicating the best result. All models were trained using BPTT.

| Dataset | Model | Package | Test accuracy [%] |
| --- | --- | --- | --- |
| MNIST | SpyLIF100+SpyLI | NeuroTorch | **96.37** |
|  | SpyLIF100+SpyLI | SpyTorch<sup>50</sup> | 94.60 |
|  | LIF100+FC | Norse<sup>49</sup> | 95.35 |
| Fashion-MNIST | SpyLIF100+SpyLI | NeuroTorch | **85.58** |
|  | SpyLIF100+SpyLI | SpyTorch<sup>50</sup> | 83.40 |
|  | LIF100+FC | Norse<sup>49</sup> | 82.97 |
| Heidelberg | SpyLIF200+SpyLI | NeuroTorch | 74.16 |
|  | SpyLIF200+SpyLI | SpyTorch<sup>50</sup> | 66.70 |
|  | LIF200+FC | Norse<sup>49</sup> | **75.68** |

**Table 4.1.** Summary of the variables of the e-prop algorithm.

| Variable | Size | Description |
| --- | --- | --- |
| $x$ | $(B, f_{in})$ | Input of the network. |
| $\hat{y}$ | $(B, f_{out})$ | Output of the network. |
| $y$ | $(B, f_{out})$ | Desired output of the network. |
| $\varepsilon$ | $(B, f_{out})$ | Prediction error. |
| $z_\ell$ | $(B, N_{\ell,out})$ | Output of the layer $\ell$. |
| $\hat{\Theta}_{\ell,k}$ | $(N_{\ell,k,in}, N_{\ell,k,out})$ | Parameter $k$ of the layer $\ell$. |
| $R_{\ell,k}$ | $(f_{out}, N_{\ell,k,out})$ | Feedback matrix of the parameter $k$ of the layer $\ell$. |
| $L_{\ell,k}$ | $(1, N_{\ell,k,out})$ | Learning signal of the parameter $k$ of the layer $\ell$. |
| $e_{\ell,k}$ | $(N_{\ell,k,in}, N_{\ell,k,out})$ | Eligible trace of the parameter $k$ of the layer $\ell$. |

### 2.2 Neuronal activity prediction

To investigate the applicability of the package to model neuronal activity data, we collected a neuronal activity dataset from the ventral habenula of a larval zebrafish. The dataset comprises recordings from 522 neurons, obtained through two-photon imaging on 6 dpf larval zebrafish expressing the genetically-encoded calcium indicator GCaMP6s. The neuronal activity of head-restrained larvae in agarose was captured under darkflash stimuli. We then extracted the time series describing the time evolution of each neuron’s activity. Subsequently, we implemented two network models in NeuroTorch, one with fining-rate (WC) dynamics and the other with spiking (SpyLIF-LPF) dynamics, each containing exactly 522 units. We trained these models with the learning algorithm e-prop to reproduce the activity of all recorded neurons. Once trained, a model can predict the activity of each neuron at all time *t* > 0 given the initial activity of all neurons at time *t* = 0. Figure 2.1 showcases heatmaps illustrating the activity of the zebrafish, along with corresponding predictions based on WC and SpyLIF-LPF dynamics. Neuronal measurements were categorised into 13 clusters using the *k*-means algorithm, effectively highlighting the distinct activity patterns observed in the data.

**Figure 2.1.**
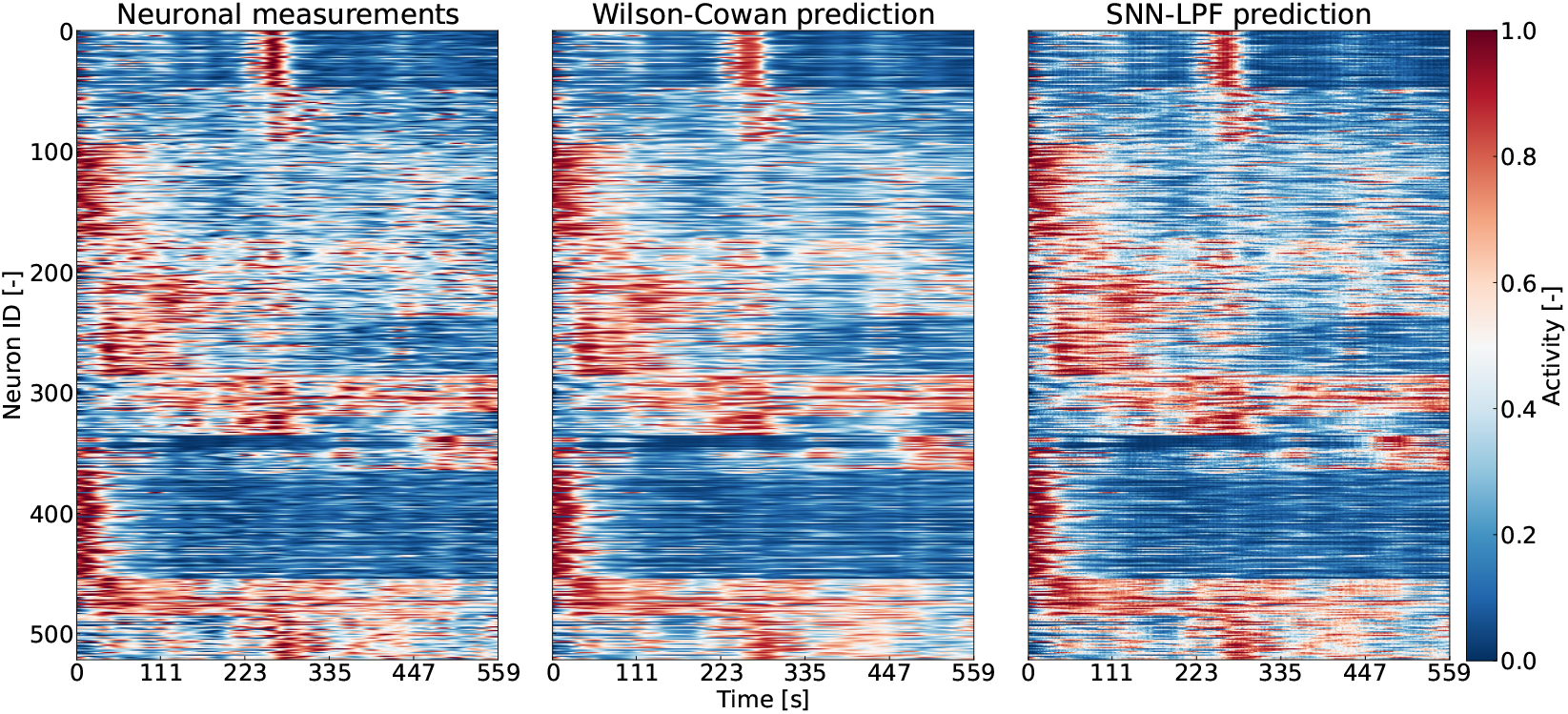
Heatmaps describing the normalized neuronal activity in the ventral habenula of a larval zebrafish and corresponding predictions based on Wilson-Cowan and SpyLIF-LPF dynamics. The neuronal activity measurements were grouped into 13 neuron clusters by the *k*-means algorithm to highlight the different activity patterns.

Figure 2.2 shows neuronal measurements and corresponding predictions obtained using NeuroTorch. The real data describe the time evolution of neurons’ activity in the ventral habenula, smoothed using Gaussian smoothing with *σ* = 10. The figure includes two metrics: the macro pVar (equation 4.18), which quantifies the overall similarity between the real and predicted series, and the average and standard deviation of the micro pVar (equation 4.19), which assesses the prediction accuracy at the individual neuron level. Panel **(a)** represents all original time series and WC predictions as two distinct trajectories projected into a 2D UMAP space^72, 73^, confirming the overall good performance with a macro pVar of 0.96 and a micro pVar of 0.95±0.03. Panel **(b)** compares the real series with a typical prediction obtained by NeuroTorch on the dataset. Panels **(c)** and **(d)** present the corresponding results obtained with SpyLIF-LPF dynamics. The overall prediction demonstrates a macro pVar of 0.97 with a micro pVar of 0.96±0.01. These results highlight the accuracy and reliability of NeuroTorch in predicting time series of neuronal activity in the ventral habenula. Considering that the maximum pVar in this situation is 1.0, both WC and SpyLIF-LPF predictions are very close to the ground truth. The UMAP projection in 2D dimensions serves precisely to visualize how the entire modeled population behaves compared to the ground truth, providing a comprehensive overview of the prediction performance. Furthermore the panels with the UMAP projection suggest that not only have individual neuron activities been well predicted, but the entire time series is consistent. Overall, our library offers numerous tools to model neuronal activity recordings from a standard and widespread imaging modality. From trained network models, connectivity can be studied and deconstructed numerically, a process we describe in the following section.

**Figure 2.2.**
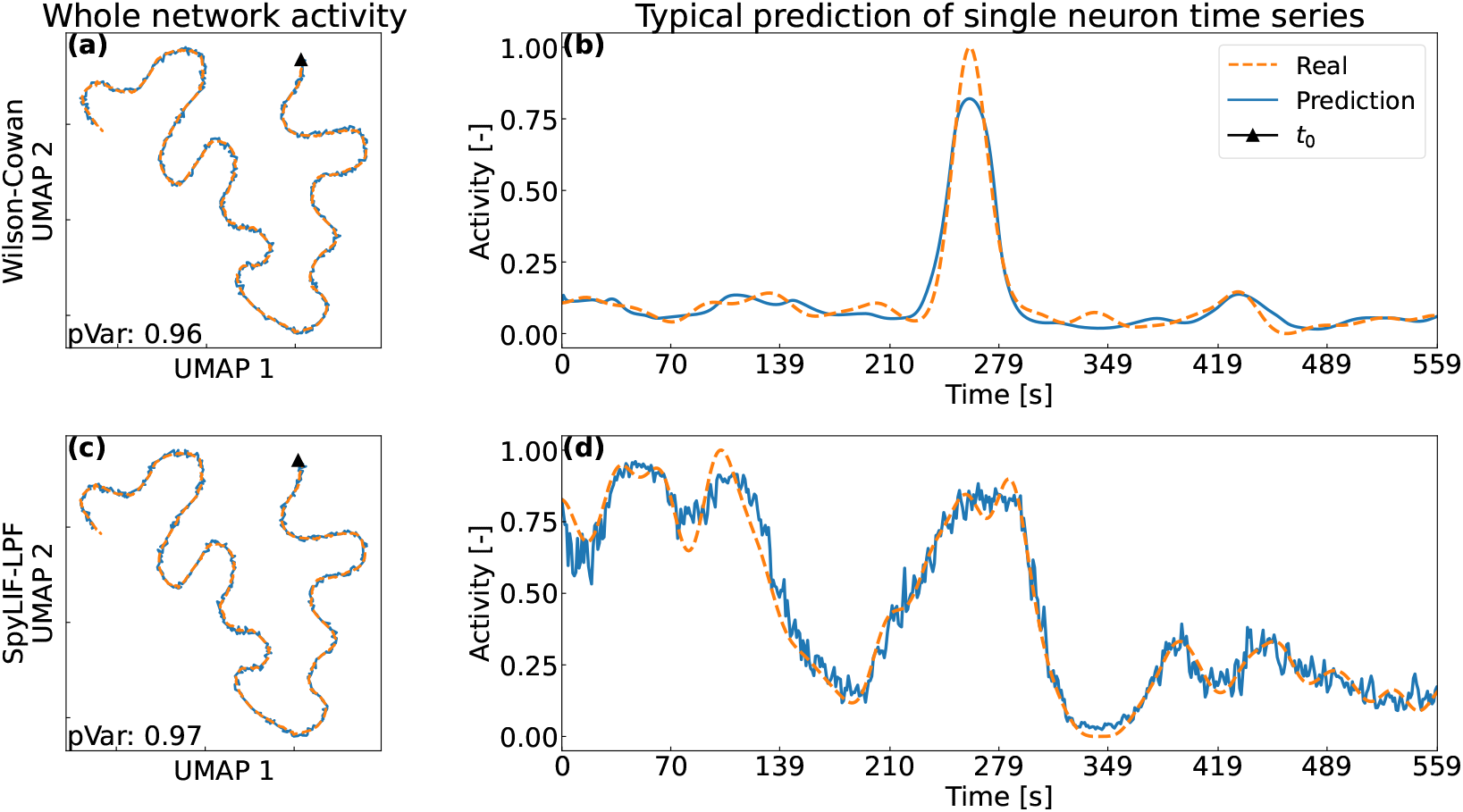
Comparison of neuronal activity measurements and corresponding predictions obtained using NeuroTorch. The real data describe the time evolution of single-neuron activity in the ventral habenula of a larval zebrafish displayed in Figure 2.1, Gaussian-smoothed with *σ* = 10 along the time axis. The metrics used are respectively the macro pVar (equation 4.18) and the average and standard deviation of the micro pVar (equation 4.19). **(a)** All smoothed original time series and Wilson-Cowan predictions projected as trajectories in a 2D UMAP space^72^, confirming the overall good performance, where the triangle denotes the initial time step. This time course prediction has a macro pVar of 0.96 with a micro pVar of 0.95±0.03. **(b)** Real series vs. typical prediction obtained by NeuroTorch on the dataset. **(c and d)** Corresponding results obtained with a SpyLIF-LPF dynamics. The whole time course prediction has a macro pVar of 0.97 with a micro pVar of 0.96±0.01.

### 2.3 Resilience of predictions under connection removal

The animal and human brains exhibit a remarkable level of sparsity, indicating that only a small fraction of all possible connections between neurons or brain regions is observed.^74–78^ In the pursuit of developing biologically plausible neural network models, it becomes intriguing to explore methods for maximizing sparsity and creating networks that closely resemble the architecture of the brain. Building upon this line of investigation, we conducted a computational experiment aimed at determining how the choice of dynamics and the application of Dale’s law could influence the capacity of a network model to maintain realistic neuronal activity while increasing its sparsity. We thus studied the resilience of different network models trained on calcium imaging data when subjected to connection loss.

We selected four types of recurrent network models, each having 522 units, as follows: we either chose the SpyLIF-LPF or the WC neuronal dynamics and we imposed or not the network connections to comply with Dale’s principle. When the Dale’s principle is enforced, every outgoing synaptic connections of a neuron are either positive or negative. All models were first trained with e-prop to adequately predict the neuronal activity data (see previous section). We then submitted the models to two distinct connection ablation strategies: (1) hierarchical connection ablations, where the connection with the smallest weight (in absolute value) is successively removed; (2) random connection ablations, where at each step, a randomly selected connection is removed, disregarding its weight. To assess the resilience of the models under connection removal, we computed the pVar each time a connection was removed and then compared it to the original pVar to obtain a performance ratio.

Figure 2.3 summarizes the results of this analysis. Panel **(a)** shows that in the case of hierarchical connection ablations, the performance-ratio curve for WC+Dale initially reaches a sparsity of approximately 0.2 without significant change before rapidly decreasing. In contrast, the corresponding curve for SpyLIF-LPF+Dale reaches a higher sparsity level of around 0.4 before experiencing a significant decline. These findings suggest that the SpyLIF-LPF dynamics tolerates a higher level of sparsity compared to the WC dynamics. Moving on to Panel **(b)**, which focuses on random connection ablations, all performance-ratio curves demonstrate a rapid decline with respect to sparsity. As for the hierarchical connection ablation, the WC+Dale curve remains higher than the WC curve, suggesting potential advantages of incorporating Dale’s law. Interestingly, the SpyLIF-LPF curve not subjected to Dale’s law shows a slower initial decline than the corresponding curve subjected to Dale’s law, but the extent to which Dale’s law contributes to this particular dynamics in the context of random connection ablations remains uncertain. Finally, Panel **(c)** shows the Area Under the Curve (AUC), which measures the average performance ratio over sparsity, for each curve in Panel **(a)**, while Panel **(d)** presents the AUC for each curve in Panel **(b)**. Analyzing Panel **(c)**, it is clear that Dale’s law enhances the resilience of network models based on both types of dynamics against hierarchical ablation. However, Panel **(d)** indicates that the distributions of AUCs for the SpyLIF-LPF dynamics are not statistically significantly different, making it challenging to draw definitive conclusions. Conversely, the distributions of AUCs for the WC dynamics in Panel **(d)** show that Dale’s law contributes to their overall resilience against this type of ablation. Therefore, based on these observations, we conclude that Dale’s law and the spiking dynamics have a positive impact on the resilience of network models trained on experimental data, especially in the context of hierarchical connection ablation.

**Figure 2.3.**
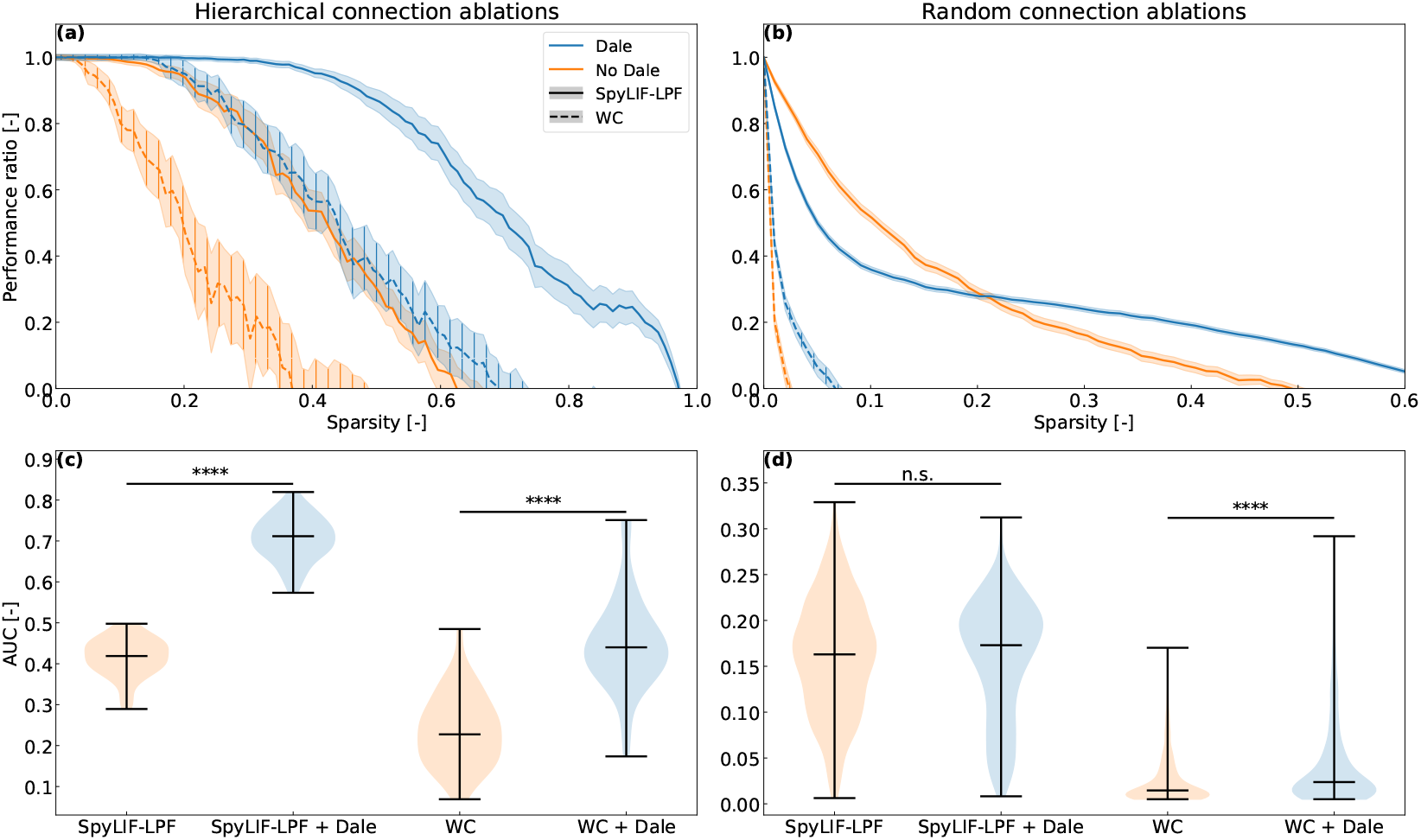
Resilience analysis of the network models based on the SpyLIF-LPF and the WC dynamics, with and without the enforced Dale law, all trained with e-prop. **(a)** Hierarchical connection ablations: at each step, the connection with the smallest weight in absolute value is removed from the network. **(b)** Random connection ablations: random connections are successively removed from the network. Here 32 random seeds by model are used to choose the ablated weights which produces a distribution of performance ratios for each sparsity value. Each solid or dashed line follows the mean of the corresponding distribution while each shaded region around a line represents a 95% confidence interval. **(c, d)** Area under the curve (AUC) for each curve in (a) and (b) respectively, where **** indicates that the p-value is less than 10^−4^ while n.s. stands for non statistically significant.

## 3 Discussion

The accuracy results of 96.37%, 85.58% and 74.16% shown in table 2.1 for all the benchmark datasets respectively demonstrate that the training pipeline and the dynamics implemented in NeuroTorch work as intended. Indeed, these results are superior to SpyTorch and Norse for image classification and slightly inferior for audio classification. It should be noted that it is possible to obtain better results with NeuroTorch than those presented by increasing the number of neurons used or by changing the dynamics used. This configuration was chosen to ensure a fair comparison among different packages.

In 2020, Bellec et al.^35^ introduced the eligibility-propagation algorithm (e-prop), a biologically plausible, online learning algorithm that approaches the performance of backpropagation through time (BPTT). However, a major limitation of existing implementations is that the computation of the eligibility trace is hard-coded, requiring the derivative of the layer’s output with respect to each parameter to be computed manually, as described by equation 4.14. In other words, the derivative of the output 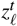 with respect to each parameter must be computed analytically and implemented manually, a process that can be tedious and daunting for newcomers. NeuroTorch provides a simple and efficient solution to this limitation. NeuroTorch computes the eligibility trace by exploiting the computation power of the python module PyTorch. It leverages PyTorch’s automatic differentiation engine to compute the eligibility trace for *any* dynamic, eliminating the need for analytical derivation or manual implementation. Hence, NeuroTorch users can easily add custom dynamics and apply the e-prop algorithm with just a few lines of code. Our implementation significantly accelerates and simplifies the use of the e-prop algorithm.

The e-prop algorithm is well suited for the reproduction of time series. Indeed, the results presented in figure 2.2 reveal that NeuroTorch’s implementation of e-prop can optimize a model that reproduces with great precision and fidelity the neuronal activity both on a local and global scale. The trajectory of the predicted time series in the UMAP space fits well the prediction of the real time series. The match between both the predicted and real trajectory in UMAP shows that, *in general*, the time series is well reproduced. This is also confirmed by the precision of our prediction. A relative precision of 3.2% and 1.0% is achieved for the prediction with the Wilson-Cowan and SNN dynamics respectively. The small uncertainty means that most neurons are well fitted. This conclusion is important because a high macro pVar can be achieved if the trainer fits well the neuron with poor activity and fits badly the neuron that are active. This is especially true for a dataset where only a few neurons are highly active. Hence, the use of the pVar (instead of the MSE) allows us to not only fit the mostly inactive neurons, but also to fit those who are highly active. Furthermore, the high quality of the prediction for each individual neuron can be confirmed by comparing the heatmaps of the predicted time series with the real heatmap. In fact, the prediction of a typical neuron is shown in the figure 2.2 where the high micro pVar indicates that the variance is preserved.

Beyond prediction accuracy, the exceptional sparsity of biological neuronal networks raises important questions about their resilience to disruptions. NeuroTorch’s computational models allow for the systematic exploration of how sparsity and hierarchical connection ablation affect network performance. In biological systems, connectivity is exceptionally sparse, with densities as low as one in ten million^79^. By simulating such conditions, NeuroTorch enables testing of resilience under biologically inspired constraints, such as Dale’s law. As shown in figure 2.3, networks trained with Dale’s law maintain performance at higher sparsity levels, particularly under hierarchical ablation. This highlights the role of biologically plausible principles in enhancing the robustness of both spiking neural networks and Wilson-Cowan dynamics.

The study highlights NeuroTorch’s ability to combine accurate performance with biologically inspired principles to advance the analysis of neuronal networks. By enabling the exploration of resilience, sparsity, and hierarchical ablation in computational models, NeuroTorch facilitates the integration of biological realism and computational efficiency. These findings underscore its potential to contribute to a deeper understanding of the relationship between network structure and function while providing a robust platform for further research.

## 4 Methods

### 4.1 Neuronal network models

To perform data prediction using artificial intelligence, specific network architecture, neuronal dynamics and learning algorithms must be specified. The dynamics determine how artificial neurons communicate and process information in the network, while learning algorithms update network parameters, including connection weights, to minimize a loss function. In this section, we present the dynamics and the learning algorithms used in this paper for validation, including the prediction of experimental neuronal activity. Additional methods are also available in NeuroTorch^58^; please refer to the library’s documentation for a comprehensive list of implemented modules.

#### Network architecture

Each network contains a fixed number *N* of units, here interpreted as neurons. The network is moreover divided into *L* sub-networks, called layers. The *ℓ*-th layer contains *N*_*ℓ*_ neurons, where *ℓ* ∈ {1, …, *L*}. Hence, *N*_1_ + *N*_2_ + … + *N*_*L*_ = *N*. The set of connections going from layer *ℓ* to layer *m* is encoded into a *N*_*ℓ*_ × *N*_*m*_ matrix *W*^*ℓm*^, whose (*i, j*) element is a real number that gives the weight of the connection going from *i*-th neuron in layer *ℓ* to the *j*-neuron in layer *m*, denoted as *I* → *j*. By convention, there is:

- an inhibitory connection from *i* to *j* if 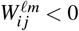,
- no connection at all from *i* to *j* if 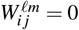,
- excitatory connection from *i* to *j* if 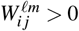.

Inside layer *ℓ*, the neurons may be connected among themselves, forming a recurrent network. The latter is described by the matrix *W*^rec^ = *W*^*ℓℓ*^, with the element 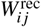 corresponding to the weight of the connection from *i* to *j*, where *i, j* ∈ {1, 2, …, *N*_*ℓ*_}. When focusing on the connections arriving at layer *ℓ* and coming from layer *k*, we write 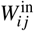 to denote the weight of the connection coming from neuron *i* in layer *k* to neuron *j* in layer *ℓ*.

#### Neuronal dynamics

##### Leaky-integrate-and-fire

The leaky-integrate-and-fire (LIF) is a classical model in neuroscience^80^ that describes a neuron’s activity in terms of its membrane potential and incoming synaptic currents. Although the shape of the membrane potential is not realistic^81^, the model includes essential features of neuronal activity like: integration of the synaptic inputs, threshold-based firing, resetting of membrane potential after firing, and decay of the membrane potential over time. In its networked and time-discretized version^35, 50^, the LIF model is determined by the recursion equation

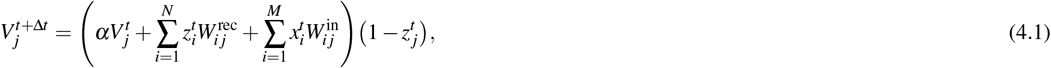

where *N* designates the number of neurons of the layer, 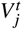 denotes the membrane potential of neuron *j* at time step *t*, Δ*t* denotes the time step of the Euler integration, 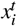 denotes the input coming from neuron *i* at time *t* in another layer of size *M*. In the eventuality that 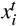 is a function of 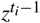 then *W*_*in*_ is considered recurrent. Moreover, the parameter *α* is a decay constant that can be written as 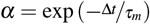, with *τ*_*m*_ being the decay time constant of the membrane potential, which is usually 20 ms. Finally, 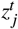 denotes the output of neuron *j* at time *t*, which is defined as

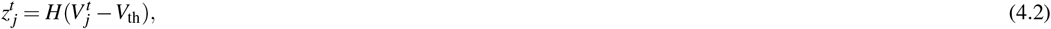

where *V*_th_ denotes the activation threshold of the neuron and *H* is the Heaviside step function satisfying *H*(*x*) = 1 when *x* ≥ 0 and *H*(*x*) = 0 otherwise.

##### Leaky-integrate with explicit synaptic current

We call leaky-integrate with explicit synaptic current (SpyLI), a dynamics inspired by Neftci’s SNN equations^50^ that integrates the input of the layer without generating spikes. Although rarely used as an input or hidden layer in SNNs, it is commonly employed at the network’s output due to its continuous activity. SpyLI is represented by two differential equations for the membrane potential and synaptic current, serving as a more complex variant of the LI dynamic introduced by Bellec et al.^35^. Using Euler integration, the synaptic current is updated by the recursion equation

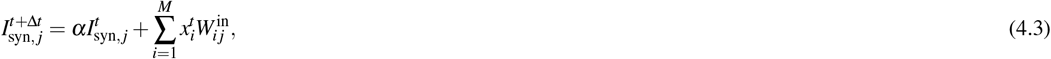

while the the synaptic potential is updated by

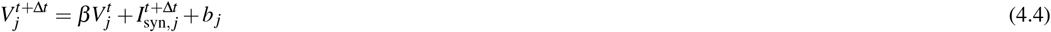

Where 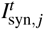 denotes the synaptic current of neuron *j* at time step *t*, 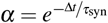 is the synaptic decay constant, 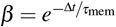 is the membrane decay constant, and *b*_*j*_ the bias weight of the layer associated with neuron *j*.

##### Leaky-integrate-and-fire with explicit synaptic current

The leaky-integrate-and-fire with explicit synaptic current (SpyLIF) dynamic inspired by Neftci’s SNN equations^50^ is a more complex variant of the LIF dynamics in Eq. (4.1) that contains two differential equations like the previously introduced SpyLI dynamics. The equation

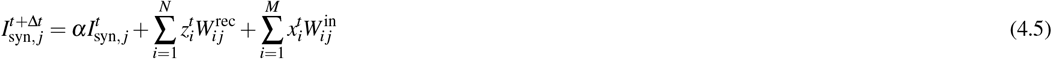

updates of the synaptic current with Euler integration while the equation

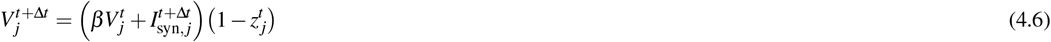

updates of the synaptic potential. As for the LIF dynamics, the output of neuron *j* at time *t*, denoted 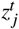, is defined in Eq. (4.2).

##### Spiking neural network - low pass filter

In some applications, such as the e-prop learning algorithm (see below), a smooth version of the spiking dynamics is required. The smoothing is achieved by the low pass filter (LPF) used by Bellec. et al^35, 82^, which modifies the vector *x*^*t*^ of neuron activity at time *t* as

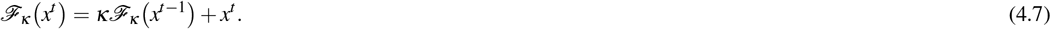

where the decay rate *κ* is set to 0.001 by default in NeuroTorch. However, this decay rate can be changed to zero if no filter is required. Every model of the spiking dynamics presented in this paper includes an LPF version in which the output spikes of the neural network are smoothed.

##### Wilson-Cowan

Neurobiologists often measure calcium activity as an approximation of neuronal activity. By its continuous nature, the Wilson-Cowan (WC) dynamics^17, 65–68^ is well-suited to model this type of signal. It describes the time evolution of spiking rates or activation probabilities of neuronal units (either neuron populations or single cells) in terms of their interactions occurring along the network connections. In this paper, we use the time-discretized form of the Wilson-Cowan model:

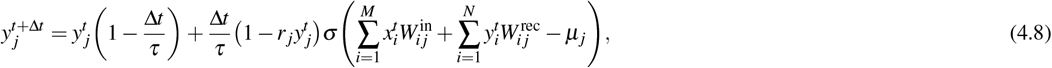

Where 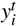 denotes the activity of unit *j* at time *t, τ* designates the rate of decrease of the activity, *r*_*j*_ is the transition rate of unit *j*. This parameter represents the time a unit needs to “recover” before becoming “sensitive” to activation. The symbol *σ* stands for the sigmoid function while *µ*_*j*_ designates the activation threshold of unit *j*.

##### Learning algorithm

The eligibility trace forward propagation (e-prop) learning algorithm^35, 82^ performs weight updates using only local information, meaning that a connection’s weight at time *t* + Δ*t* changes as a function of the activity of the adjacent neurons at time *t*, without considering the activity of other neurons in the network. This algorithm is biologically plausible since the rules that regulate the changes of synaptic connections of biological neurons only involve information that is local in time and space^83^.

The implementation of e-prop in NeuroTorch offers flexibility as it can be applied to all types of layers. Additionally, the level of locality in the implementation can be adjusted according to the user’s choice. Initially, lets consider a network with *L* layers containing the parameters 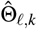 where *ℓ* is the index of the layer and *k* is the index of the parameter in this layer. The layer indexes *ℓ* ∈ [0, *L* − 1] begin at zero which is the index of the input layer. Each parameter can be a scalar, a vector, a matrix or a tensor. In the following equations, the indices *t* indicates the current time step. The input and output tensors of the network denotes respectively *x* and *ŷ* while the desired output tensor *y*. For the layer-specific inputs and outputs of the model, these are denoted by the quantity *z* with the appropriate subscript *ℓ* if necessary. The prediction error is thus defined as

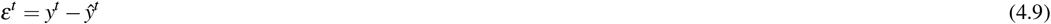

where the symbols *ŷ* and *y* are of size (*B, f*_*out*_), with *B* being the number of data in the current minibatch (a subset of the dataset) and *f*_*out*_ the number of output values. Therefore the quantity *ε* is of size (*B, f*_*out*_). Figure 4.1 provides a simplified representation of the e-prop algorithm as implemented in NeuroTorch, while Table 4.1 summarizes the variables utilized in the algorithm. The loss of the network is determined by the user-defined function

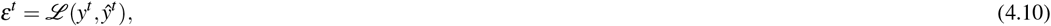

where *ε*^*t*^ is a scalar. Then, the main goal of the current algorithm is to minimize

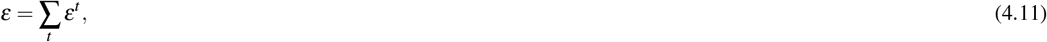

which represents the overall loss over time. To achieve this in an online and biological way, the network needs to be updated with local information. For more clarity, we divide the update of the network parameters into two steps: the update of the parameters of the output layer of the network and the update of the other parameters.

**Figure 4.1.**
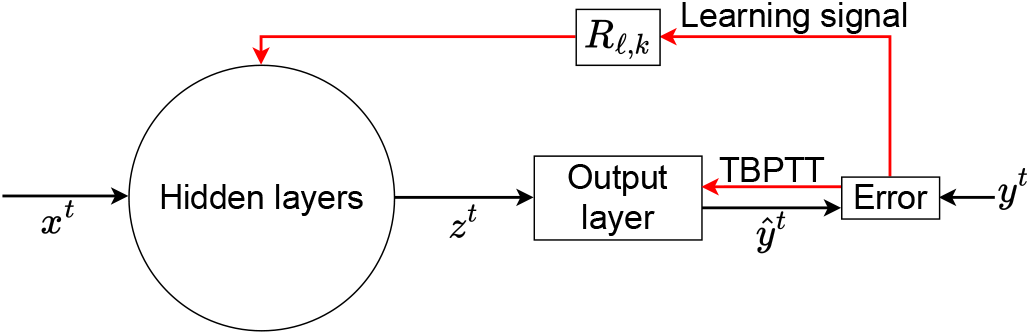
Representation of the eligibility-propagation algorithm. The hidden layers contain the desired dynamics while the output layer is generally a fully-connected linear layer. The parameters of the output layers are trained with TBPTT while those of the hidden layer are trained by computing a learning signal through a random matrix specific for each parameters.

#### Output parameters

The update of the output parameters is similar to an update by TBPTT^84^. So the gradient of these parameters will be determined by the recursive equation

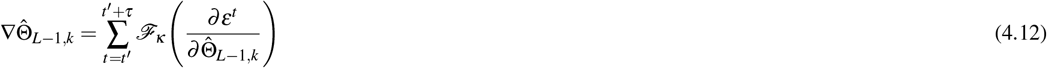

where the quantity *τ* is the number of time steps that the gradient is accumulated. To be more local in time and also more biologically accurate, *τ* must be as close to one as possible. However, contrary to Bellec’s work^35^, in which *τ* is equal to one, NeuroTorch allows the user to determine it’s own *τ*. Hence, *τ* is now simply an hyper-parameter that can be adjusted by the user. In addition, the low pass filter ℱ_*κ*_ mentioned in Equation 4.12 is defined in Eq. (4.7).

#### Hidden parameters

The update of synaptic weights according to e-prop is done by approximating the gradient of the backpropagation throught time (BPTT)^35, 59^ as the set of partial derivatives of the user defined prediction error *E*,

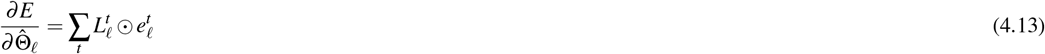

where *L*_*ℓ*_ is the learning signal and *e*_*ℓ*_ is the eligible trace. Here, the operator ⊙ is defined as the element-wise product (a.k.a. Hadamard product). This eligible trace is described by the equation

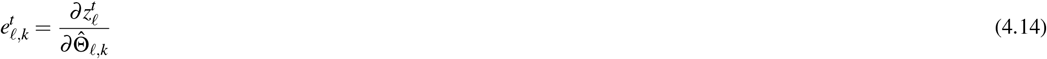

which can be simply interpreted as the partial derivative of the local output signal. The learning signal defined as

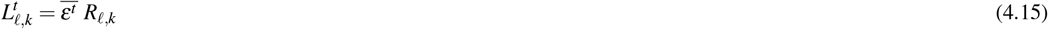

is interpreted as the error feedback from the network output with *R*_*ℓ*,*k*_ as a feedback matrix that can be initialized in several ways described in^35, 85^ with the nomenclature *B*_*jk*_. To maintain the locality of the algorithm, *R*_*ℓ*,*k*_ is often initialized as a random matrix to broadcast a random feedback to the hidden parameters which is the case in this work. Using these last expressions, the gradient of the weights 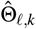 is equal to

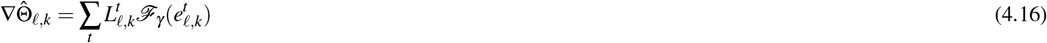

which is used in a gradient descent method to optimize the weights.

In the NeuroTorch implementation of the e-prop algorithm, it is possible to clip some variables to stabilize the training. Among these variables, there are the gradients of the parameters, the eligibility trace, the learning signal and the feedback weights.

##### Application of Dale’s law

To ensure mathematical viability and biological realism in predicting neuronal time series, NeuroTorch implements the Dale law. By enforcing this well-known law discovered by Sir Henry Dale in 1935, which states that a neuron can release only one type of neurotransmitter for all its synapses^86^, NeuroTorch restricts the solution space and converges towards a biologically plausible neuronal connectome. This is achieved by decomposing the connectivity matrix into a sign vector *S* and a weight matrix, with the sign vector representing the excitatory or inhibitory property of each neuron and the weight matrix squared to ensure positive weights. Both the vector and the square root of the weight matrix are trained by the learning algorithms, as shown the equation

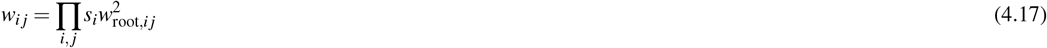

where **W**_root_ is the square root element-wise of the weight matrix.

##### Evaluation metrics

The proportion of variance metric (pVar)^62^ is a loss function used to train neural networks on times series and to evaluate their effectiveness. This metric is bound between [ − ∞, 1] where 1 is the optimal value. The pVar is essentially a mean square error (MSE) that is weighted by the variance of the timeseries. By doing so, we ensure that the timeseries with higher variance are well trained rather than just being averaged by the prediction. There are two variants of the metric, one is the macro pVar that evaluates the overall performance over all variables of the times series and the other is the micro pVar that evaluates the individual matches of the variables.

Let *X*_*t,i*_ be the predicted time series of variable *i* (neuron or unit) at time step *t* and *Y*_*t,i*_ the corresponding target time series. The macro P-Variance is describe by the equation

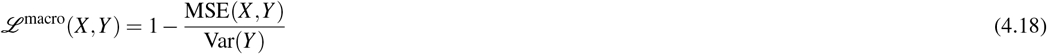

and the micro P-Variance is describe by the equation

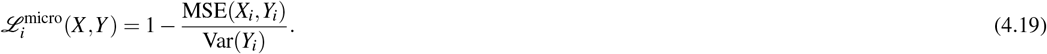

Where the macro version is a scalar, the micro one is a vector giving the P-Variance for all variables in the times series. In this case, it’s useful to show the micro pVar in form of it’s average 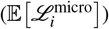 and standard deviation 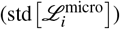 statistics.

### 4.2 Numerical experiments with NeuroTorch

#### Performance validation against benchmark datasets

The NeuroTorch performances were compared to the packages SpyTorch and Norse onto three benchmark datasets: MNIST, Fashion-MNIST and Heidelberg.

To obtain the prediction accuracies with the SpyTorch network^50^, the authors’ code was run with minimal modifications. For the MNIST dataset, the script “SpyTorchTutorial3.ipynb” was used, in which the dataset name was switched from Fashion-MNIST to MNIST, the regularization loss was removed, and the activation threshold for converting pixels to spikes was adjusted to 0.1. The other parameters remained unchanged from the version 0.4 of the repository. For the Fashion-MNIST dataset, the same script was run without any changes.

To generate results for the Norse library^49^, the script “mnist_classifiers.ipynb” from the “notebooks” subrepository was used for the MNIST and Fashion-MNIST datasets. The script came from the commit with the id “bc627fc” and was slightly modified to allow comparison with SpyTorch results. The hyperparameters used were an integration time step of *T* = 100, a learning rate of *lr* = 2*e* − 4, a hidden size of *nb*_*hidden* = 100, and a number of epochs of *epochs* = 30. The script generated results with different input encoders, but only the results from the “ConstantCurrentLIFEncoder” were used due to its higher accuracy compared to the other encoders.

The results for the Heidelberg dataset were obtained using the script “SpyTorchTutorial4.ipynb” from the version 0.4 of the SpyTorch repository^50^. To produce results for the Norse package^49^ (version 1.0.0), the model from the Mnist experiment was used, but without the encoder and with a hidden size of 200 units.

To generate the prediction accuracy results for the Heidelberg, Fashion-MNIST, and MNIST datasets with NeuroTorch, the scripts “spike_trace_classification/gen_heidelberg_results.py”, “images_classification/gen_fashion_mnist_results.py” and “images_classification/gen_mnist_results.py” of the sub-repository “NeuroTorch_PaperWithCode” were used respectively. To ensure a fair comparison, the hyperparameters were set to match those used with the other packages. The models were trained using the Backpropagation Through Time (BPTT) learning algorithm in NeuroTorch.

#### Experimental acquisition of a neuronal activity dataset

A neuronal activity dataset of 522 neurons from the ventral habenula of a larval zebrafish was collected. The neuronal activity data were acquired by two-photon imaging on 6 dpf larval zebrafish expressing a pan-neuronal genetically-encoded calcium indicator GCaMP6s. Using a resonant scanner and piezo-driven objective, the neurons of head-restrained larvae in agarose were recorded with a darkflash^87^ stimuli. Further details about the experimental methods, including the microscopy setup and visual stimuli, can be found in recent references^88, 89^.

#### Prediction validation with neuronal activity datasets

To demonstrate that the package can be used with time series we attempted reproducing the neuronal activity from the experimental dataset collected in the ventral habenula region of a zebrafish. Two networks were employed for comparison: one based on continuous dynamics (Wilson-Cowan) and the other on spiking dynamics (Leaky integrate and fire). The training utilized the e-prop learning algorithm to showcase its performance and flexibility. The model consisted of two layers: the input layer employing either Wilson-Cowan or SpyLIF-LPF dynamics, and the output layer comprising a linear layer with sigmoid activation. Specifically, for the SpyLIF-LPF dynamics, only the forward weights 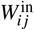 were optimized during training, while for the Wilson-Cowan dynamics, the forward weights, as well as the *τ, µ*, and *r* parameters, were optimized. If the Dale’s law is enforced in the model, the parameters 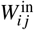 that are optimized are replaced by 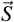 and **W**_root_ as explained in 4.1. The experiments focused on a single hidden layer per model, omitting the use of the recurrent weights 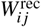. Since the model operates recurrently, meaning it uses its own predictions as inputs, the weight matrix 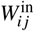is considered to be recurrent. The script to reproduce the training of these networks is “neuronal_time_series_reproduction/main.py” which can be found under the sub-repository “NeuroTorch_PaperWithCode”. To create trajectories using UMAP^72, 73^, a powerful dimensionality reduction technique, we first applied the UMAP algorithm to the target time series, projecting them into 2 dimensions, whose coordinates were denoted UMAP 1 and UMAP 2. Afterwards, we utilized the trained UMAP to perform the same transformations on the predicted time series, effectively mapping them onto UMAP 1 and UMAP 2.

### 4.3 Resilience computational experiments

The resilience of a neural network, defined as its ability to accurately predict neuronal activity even when connections are removed, was measured by evaluating its performance as a function of its sparsity. The performance was quantified as the ratio between the loss function after alteration of connectivity and the original loss function, with a performance of 1 indicating normal behaviour post-alteration. Sparsity, denoted as 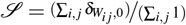, represents the proportion of unconnected pairs of neurons in the network. The resilience of trained neuronal network models was assessed using e-prop learning, with 50 trainings conducted for both SpyLIF-LPF and Wilson-Cowan dynamics, with and without the enforcement of the Dale law. Two types of ablation were performed: hierarchical connection ablation, where connections with the smallest impact on the network (i.e., smallest absolute weight values) were sequentially removed from the network setting their weight to zero in matrix *W*^in^; random connection ablation, where connections were randomly removed from the network by random selected elements in *W*^in^ and set their value to zero. Each ablation method was tested with 100 levels of sparsity, considering only the layer with the specific dynamics for ablation while keeping the output layer weights unaffected. Random ablation tests were conducted with 32 different seeds to ensure robustness of the results.

### 4.4 Design and implementation of NeuroTorch

NeuroTorch is built with a modular design and a pipeline structure. Several tutorials are available with NeuroTorch to learn how to use the package. For advanced users who want to fully exploit the pipeline features, here is an overview of its architectural characteristics.

One of the fundamental components of NeuroTorch is the sequential model, which serves as a versatile framework for sequencing a set of learning layers. As illustrated in Figure 4.2, this model adopts a “H” shape, allowing a network generalization that accommodates multiple inputs and outputs. To ensure flexibility and to remove any limitations on input and output types, NeuroTorch associates a layer and a transform to each input and output of the network. As a result of this design, the user is able to choose the number and type of layers, which can come from NeuroTorch, Norse, or any other module built with PyTorch. Since the transforms are also PyTorch modules, they can be selected by the user as well.

**Figure 4.2.**
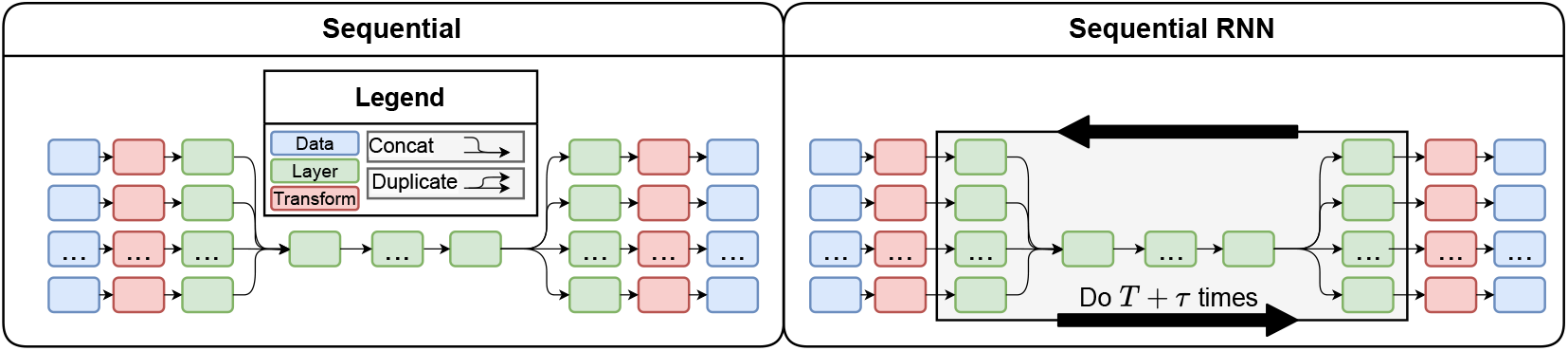
Representation of the sequential model and the recursive sequential model. *T* is defined as the time associated to the input as initial conditions. *τ* is the forecasting time used for the prediction.

Unlike other machine learning pipeline in which most of the modules are fixed, the NeuroTorch training pipeline offers flexibility and dynamic adaptability. This enables the user to modify the pipeline objects during the training process, which can, among other things, involve altering the dataset to make it progressively more challenging as the model learns. Furthermore, users have the flexibility to establish a general training configuration for a collection of experiments. By incorporating a single callback that is specific to each experiment, they can easily adapt the training process to meet the specific requirements of that particular experiment. This allows for a streamlined approach in modifying the training setup and ensuring it aligns with the unique demands of each experiment. This feature of the library relies on five objects of its implementation. First, the `trainer`, which is responsible for controlling the execution flow of the learning phase by providing the data and stopping the learning. The trainer updates the different variables of its `state` based on training results (e.g. the current iteration, the current loss value, etc). It also calls `callbacks` at appropriate times. A callback is a function that can perform multiple actions to facilitate training such as updating the datasets, the model or simply calculate metrics on the training process. The flexibility of the pipeline relies on the core of the algorithm: the learning or optimization process, which is performed by a special callback called the `learning algorithm`. This object updates the weights of the model according to a specified method (e.g. e-prop or BPTT). It also implements important events, such as the `on_optimization_begin` event that performs a round of optimization of the model parameters, and utilizes data provided by the user and managed by the `dataloader` to update the `model`’s weights. During the execution of a training, callbacks have the ability to modify the trainer’s state and behavior at runtime, such as setting a `stop_training_flag` or adjusting `n_iterations` for early stopping. The training loop with the NeuroTorch pipeline is depicted in Figure 4.3.

**Figure 4.3.**
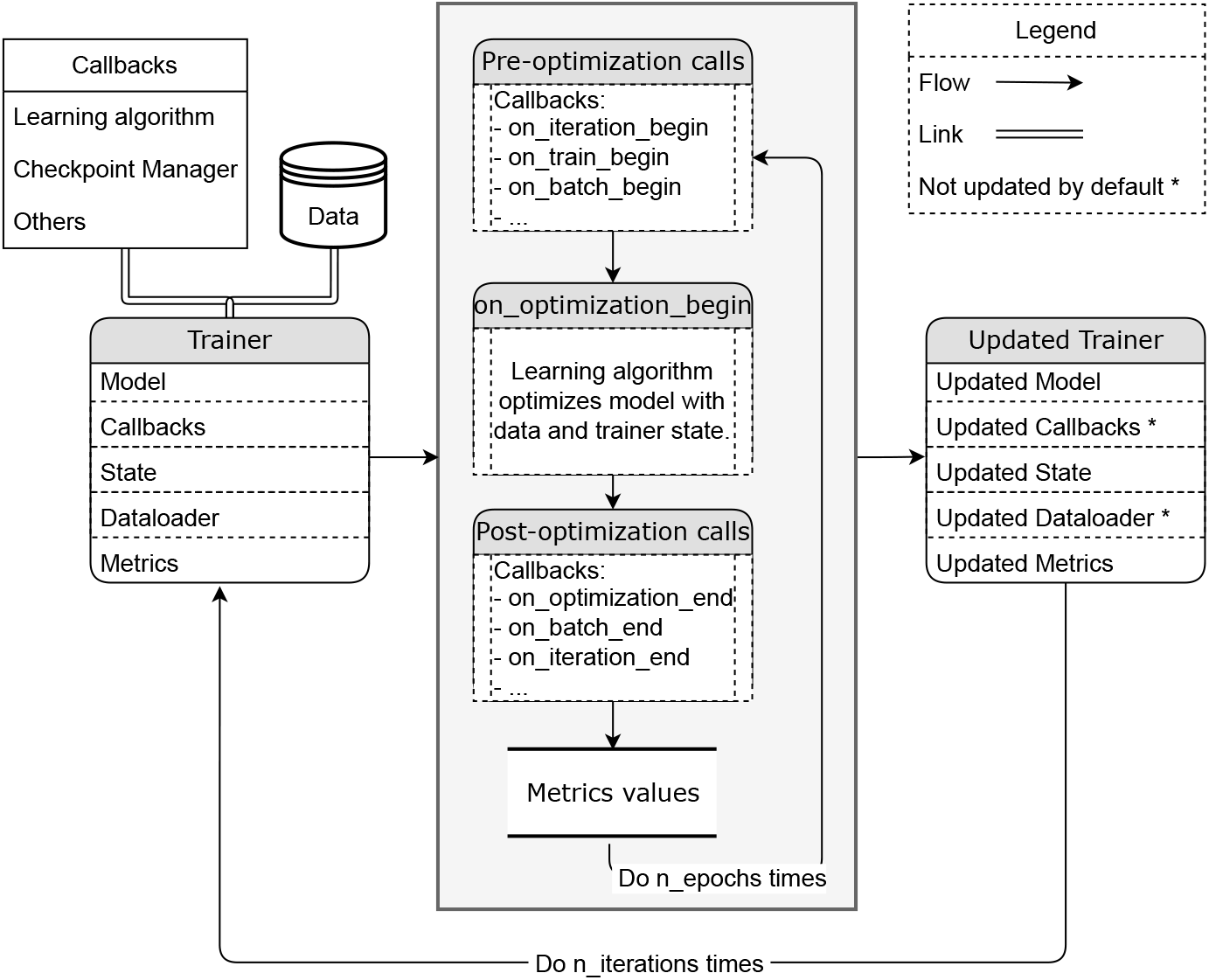
Representation of the training pipeline. The callbacks and the data provided by the user are processed using the trainer. The trainer then calls the events of the callbacks during the training. The trainer is updated accordingly.

The following definitions are necessary to understand the description of the execution of the main algorithm during the training phases. A *pass* corresponds to both the prediction of data by the model and the update of its weights. An *iteration* covers a full pass through both the training and validation datasets. It corresponds to the outer loop in Figure 4.3. An *epoch* represents a complete pass through either dataset, it corresponds to the inner loop in Figure 4.3. A *batch* signifies a forward pass through the network, and *training* and *validation* encompass full passes through their respective datasets. Just to clarify, an iteration can consist of multiple epochs (depending on the user’s choice), and each epoch can consist of multiple batches depending on the amount of data and the user’s choice of hyperparameters. After an iteration, a checkpoint state containing all the relevant information on the state of the execution of the pipeline in this instant can be saved. The user can set a callback to trigger the creation of a checkpoint state or determine a backup frequency.

The part of the algorithm executing callback events can be summarized as follows: the learning process is initiated by loading a checkpoint state and this is followed by a loop executing a specified number of iterations. Within each iteration, events occur for training and validation, including their beginnings and endings. Iterations consist of multiple epochs, which, in turn, comprise multiple batches. Each batch involves optimization-related events. The process concludes with the closing step.

## Code availability

The complete NeuroTorch package, along with its source codes, is available at https://github.com/NeuroTorch/NeuroTorch. This repository contains all the necessary information for installing NeuroTorch and accessing benchmark data. Additionally, there are comprehensive tutorials available on Google Colab that provide hands-on learning experiences with the package. You can find a tutorial on Wilson-Cowan dynamics in NeuroTorch by following this link: Wilson-Cowan tutorial. Another tutorial available is focused on MNIST: MNIST tutorial. Furthermore, there is a tutorial specifically designed for the Heidelberg dataset: Heidelberg tutorial.

## Acknowledgements

The authors are grateful to Mohamed Bahdine, whose scientific recommendations and master’s research work^90^ had a considerable impact on the elaboration of the current project. They also extend their gratitude to Antoine Légaré for his numerous comments that helped guide the project and for collecting data in Paul De Koninck’s laboratory at the CERVO Brain Research Center, and to Paul De Koninck for generously sharing this data. This work was supported by the the Natural Sciences and Engineering Research Council of Canada (P.D., S.H.), and the Sentinelle Nord program of Université Laval, funded by the Canada First Research Excellence Fund (P.D., S.H.). J.G. want to thank the UNIQUE Research Center, funded by the Fonds de recherche du Québec – Nature et technologies, for Ph.D. and M.Sc. Excellence Scholarships. The authors acknowledge Calcul Québec and Digital Research Alliance of Canada for their technical support and computing infrastructures.

## Author contributions statement

J.G., P.D., and S.V.H. designed the project. J.G. devised and coded the package and conducted the numerical experiments; A.D. assisted J.G. in his computational work. J.G. drafted the manuscript and the NeuroTorch documentation. All authors analyzed the results and reviewed the manuscript.

